# A robust and high-efficiency *Rhizobium rhizogenes* hairy root transformation platform for *Vaccinium*

**DOI:** 10.64898/2026.03.09.710070

**Authors:** Y. Kumam, F. Enciso-Rodriguez, J.H. Kim, S. Kroehler, P. Adunola, F.A. Pagliai, M. Gastelbondo, P Munoz

## Abstract

Efficient transformation remains a major constraint to functional genomics and genome editing in *Vaccinium*, where stable transformation systems are often genotype-dependent and inefficient. Here, we establish a rapid and high-efficiency *Rhizobium rhizogenes*–mediated hairy root transformation platform optimized for the genus. Using the *RUBY* visual reporter, transformation efficiency reached 46.7% in leaf explants infected with strain Ar. A4 and cultured on half-strength Woody Plant Medium, with transgenic roots visible within 16 days post-co-cultivation. Comparative evaluation of six *R. rhizogenes* strains identified Ar. A4 and ATCC15834 as consistently superior across diverse *Vaccinium* germplasm representing different taxonomic sections, achieving up to 80% efficiency in selected genotypes. While conventional regeneration from transgenic roots was not successful, overexpression of developmental regulators enabled shoot formation with 7% efficiency, demonstrating a path toward stable plant recovery. This platform delivers a rapid, genotype-flexible system for gene validation, metabolic pathway analysis, and genome editing in *Vaccinium*, substantially expanding the molecular toolkit available for perennial fruit crop research and translational breeding.

## Main Text

Efficient genetic transformation remains a critical barrier limiting functional genomics and genome editing in *Vaccinium*. The genus comprises over 450 species organized into multiple sections and ploidy levels, including several commercially important berry crops such as blueberry (*V. corymbosum*), cranberry (*V. macrocarpon*), lingonberry (*V. vitis-idaea*), and bilberry (*V. myrtillus*) (Edger *et al*., 2022). Despite its relatively recent domestication history, consumer demand for blueberry and related *Vaccinium* crops continues to expand worldwide, driven by their nutritional value and perceived health benefits (Edger *et al*., 2022). In contrast to this growing economic importance, biotechnological tools for routine plant transformation and gene validation remain underexplored. Although *Agrobacterium tumefaciens*-mediated transformation protocols have been developed, efficiencies are typically modest and strongly genotype-dependent (Wang *et al*., 2022). Genome editing has also been demonstrated in *Vaccinium* (Han *et al*., 2022); however, its broader application will require more efficient DNA delivery systems. Compared with *Agrobacterium, Rhizobium rhizogenes*-mediated transformation offers a rapid alternative to conventional stable transformation, making it an alternative for recalcitrant species and genotypes. This system induces genetically transformed roots following T-DNA transfer from Ri plasmids and has been widely adopted for gene function studies and genome editing in diverse crops (Bahramnejad *et al*., 2019). Despite its successful application in other crop species, optimization of this system for *Vaccinium* remains limited.

In this study, we evaluated the performance of *R. rhizogenes* for hairy root induction in *Vaccinium*, comparing two widely used strains, Ar. A4 and K599. Root induction was assessed on half- and full-strength Woody Plant Medium (HWPM and WPM respectively) without plant growth regulators in the blueberry cultivar ‘Albus’, with two experimental parameters into consideration, the explant type and method of wounding. Strain Ar. A4 consistently outperformed K599 across all conditions tested (Figure S1). In leaf explants cultured on HWPM, Ar. A4 achieved up to 56% hairy root induction, whereas K599 reached 16%. This 3.5-fold difference indicates that hairy root induction in *Vaccinium* is influenced by strain-host interaction and media composition. We next investigated the effect of explant and wound type in hairy root induction. Leaf explants were significantly more responsive than stem tissues. Both basal cutting and needle puncture induced roots in leaves, although basal cutting was operationally more efficient, and consistently induced higher root production (Figure S1). In contrast, only few stem explants generated hairy roots at cut surfaces. Collectively, these results demonstrate that explant type strongly influences hairy root induction efficiency.

Transformation efficiency was evaluated using the *RUBY* reporter system, which enables visual identification of transgenic roots through betalain accumulation (He *et al*., 2020), followed by confirmation of gene integration via PCR (Figure S2, Table S1). Under optimized conditions (Ar. A4, leaf explants, basal wounding, HWPM), transformation efficiency reached 46.7% at 60 days post co-cultivation (Figure 1a); in contrast, K599 achieved 11% under the same conditions. *RUBY*-expressing roots were visible as early as 16 days post co-cultivation, substantially reducing experimental timelines compared with conventional stable transformation systems (Song and Sink, 2004). A zero-inflated negative binomial mixed model confirmed that strain, explant type, and time significantly influenced transformation efficiency, whereas media strength alone was not significant (Figure 1a,b,c). These findings highlight bacterial strain, genotype, and tissue type as primary factors determining hairy root transformation.

**Figure. 1.**
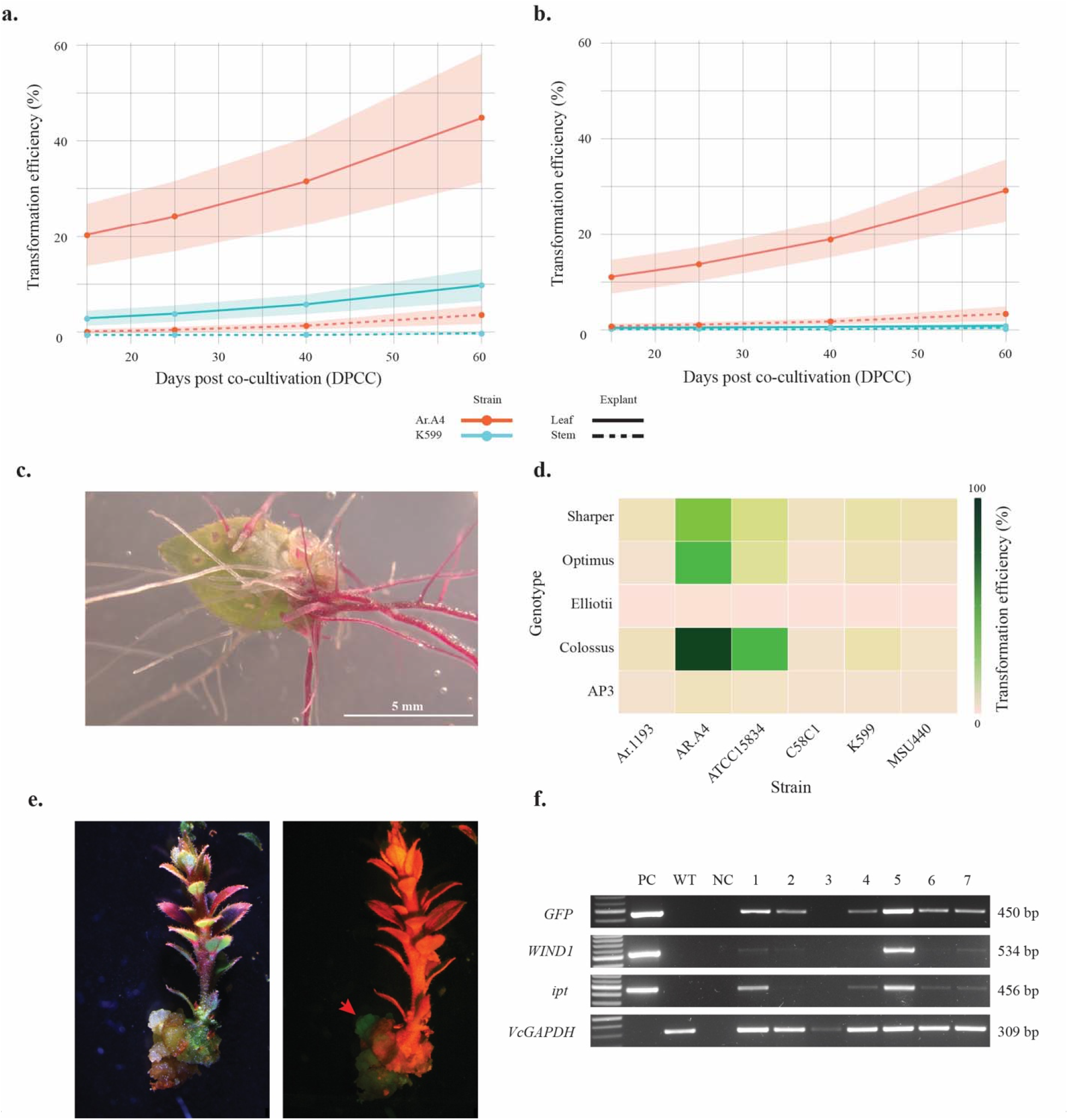
Hairy root transformation and regeneration in *Vaccinium* using *Rhizobium rhizogenes*. **a**. Transformation efficiency (%) of leaf and stem explants of ‘Albus’ co-cultivated with Ar. A4 and K599 on half-strength WPM medium (HWPM). **b**. Transformation efficiency (%) of the same genotype and explants on full-strength WPM medium. Transformation efficiency (%) was analyzed using zero□inflated beta regression models fitted separately for each DPCC strain–explant combination. **c**. Leaf explant infected with *R. rhizogenes* strain Ar. A4 showing RUBY-expressing hairy roots. **d**. Transformation efficiency of the six strains of *R. rhizogenes* Ar.1193, Ar. A4, ATCC15834, C58C1, K599, and MSU440 across different *Vaccinium* genetic backgrounds (Table S1): *V. corymbosum* (‘Sharper’, ‘Optimus’ and ‘Colossus’), *V. elliottii* (‘Elliottii’) and *V. staminium* (‘AP3’). **e**. GFP expression in the basal callus region of regenerated ‘Albus’ shoot transformed with *R. rhizogenes* strain ATCC15834 containing an overexpression plasmid with the developmental regulators *WIND1* and *ipt*. Left panel: bright light. Right panel: GFP-LP filter. Red arrow indicates GFP expression. **f**. Molecular confirmation of the regenerated shoots using PCR for the genes *WIND1, ipt*, and *GFP* present in the overexpression vector and the constitutive expressed *Glyceraldehyde-3-phosphate dehydrogenase* (*VcGAPDH*) gene as control (Table S2). PC: *RUBY*-containing plasmid, WT: Wild type ‘Albus’, NC: Water control, and 1 to 7 are regenerated shoots.

Given the pronounced strain-dependent differences observed, we further optimized the system by evaluating six *R. rhizogenes* strains under multiple antibiotic regimes to control bacterial overgrowth while maintaining high transformation efficiency. Consistent with initial results, K599 exhibited 25% bacterial overgrowth under treatment T1 (Table S2), which was effectively suppressed by the addition of timentin (Figure S3). For Ar. A4, bacterial overgrowth was well controlled under all treatments, except T4 and the no-antibiotic control (T11), achieving highest transformation efficiencies under T1 (75%) and T6 (65%) (Figure S3). ATCC15834 showed a similar pattern, reaching up to 62.5% transformation efficiency under T9, and 55% under T5 and T6. In contrast, Ar.1193 exhibited low transformation efficiency (<10%) across all treatments. Although bacterial overgrowth in C58C1 and MSU440 was effectively controlled under most treatments, C58C1 failed to generate transgenic roots and MSU440 displayed low efficiency (<10%) (Figure S3). Overall, Ar. A4 and ATCC15834 consistently outperformed the other strains, highlighting the importance of host-strain compatibility, as also reported in other species such as strawberry (Yan *et al*., 2023).

Based on these findings, we next evaluated the robustness of the system across additional *Vaccinium* genotypes representing two taxonomic sections (Table S3). Ar. A4 successfully induced *RUBY*-expressing roots in all tested genotypes, with transformation efficiencies ranging from 1.9% in ‘Elliottii’ to 85% in ‘Colossus’, while C58C1 performed the least, with the highest transformation efficiency in Albus at 8% (Figure 1d, Table S4). The substantial variation observed among genotypes and performance of the strains highlights the influence of host-strain interactions on transformation efficiency, while also demonstrating the broad applicability of Ar. A4 across diverse *Vaccinium* germplasm.

Plant regeneration following hairy root transformation was first evaluated using media supplemented with exogenous plant growth regulators. In this initial approach, *RUBY*-expressing roots were transferred to callus induction medium supplemented with different concentrations and combinations of auxins and cytokinins in HWPM media (data not shown). Although transgenic hairy roots readily dedifferentiated into callus and produced globular somatic embryos, indicating substantial developmental plasticity, no shoot regeneration was observed (Figure S4). Given the absence of shoot formation, a second strategy was implemented to overcome potential regeneration constraints. This approach relied on the overexpression of the developmental regulators *WOUND INDUCED DEDIFFERENTIATION 1* (WIND1) and *ISOPENTENYL TRANSFERASE* (ipt) (Kshetry *et al*., 2025) and included GFP as a visual marker to enable real-time screening of transformed tissues (vector kindly gifted by Dr. Patil, Texas Tech University). Using this strategy, shoot regeneration was achieved in 7% of ‘Albus’ explants under the standardized transformation conditions. Molecular validation and GFP fluorescence analysis confirmed transgene integration in most regenerated shoots (Figure 1e, f, S5).

Shoot formation was observed starting at 40 days post co-cultivation on HWPM without any plant growth regulators. While a high transformation efficiency of 76% was observed, as indicated by GFP expression in the callus, a marked reduction in root formation was observed following the use of developmental regulators, with only 10% of explants producing roots. Similar studies using developmental regulator-mediated strategies have demonstrated enhanced regeneration efficiency in several plant species using *Agrobacterium*-mediated transformation (Kshetry *et al*., 2025). Our findings extend this approach to *R. rhizogenes*–mediated systems, demonstrating its applicability in *Vaccinium* and supporting its potential for establishing stable transformation pipelines in the genus.

Although GFP expression was clearly detected in callus tissue (Figure 1e), fluorescence was not observed in the aerial tissues during shoot emergence (Figure 1e). This reduction in detectable fluorescence may be associated with red-chlorophyll accumulation in green tissues, which can mask GFP signal (Chung *et al*., 2000). Consistent with this explanation, GFP expression was evident in promeristems, but not in mature pigmented tissue (Figure S4).

In summary, we establish a rapid and efficient *R. rhizogenes*-mediated hairy root transformation system for *Vaccinium*. Strain Ar. A4 combined with leaf explants on half-strength WPM provides optimal performance, with visible *RUBY* expression within 16 days. The platform enables transformation across different genetic backgrounds and supports shoot regeneration through developmental regulators. This system expands the molecular toolkit available for enabling rapid gene validation, metabolic pathway analysis, and genome editing in economically important *Vaccinium* species, offering a practical route to accelerate molecular breeding efforts across the genus.

## Data availability

The datasets generated during and/or analyzed during the current study are available from the corresponding author on reasonable request.

## Acknowledgements

We are grateful to Shane Teachworth for copy-editing the manuscript.

## Funding

This work was supported by royalties from the University of Florida Blueberry Breeding program.

## Contributions

FER and PM formulated the idea of the manuscript. FER, SK, JHK, YK and MG wrote, reviewed, and edited the manuscript. SK, JHK, and YK performed the experiments. FER created the figures. MG and PA performed data analyses. PM and FAP reviewed and edited the manuscript. All authors read and approved the manuscript.

## Ethics declarations

### Conflict of interest

The authors report no conflicts of interest in this work and have nothing to disclose.

## Notes

### Competing Interest Statement

The authors have declared no competing interest.

## References

Bahramnejad, B., Naji, M., Bose, R., and Jha, S. (2019) A critical review on use of Agrobacterium rhizogenes and their associated binary vectors for plant transformation. Biotechnol. Adv., 37.

Chung, B.-C., Kim, J.-K., Nahm, B.H., and Lee, C.-H. (2000) In Planta Visual Monitoring of Green Fluorescent Protein in Transgenic Rice Plants. Mol. Cells, 10, 411–414.

Edger, P.P., Iorizzo, M., Bassil, N. V, Benevenuto, J., Ferrão, L.F. V, Giongo, L., et al. (2022) There and back again; historical perspective and future directions for Vaccinium breeding and research studies. Hortic. Res., 9, uhac083.

Han, Xiaoyan, Yang, Y., Han, Xue, Ryner, J.T., Ahmed, E.A.H., Qi, Y., et al. (2022) CRISPR Cas9- and Cas12a-mediated gusA editing in transgenic blueberry. Plant Cell Tissue Organ Cult., 148, 217–229.

He, Y., Zhang, T., Sun, H., Zhan, H., and Zhao, Y. (2020) A reporter for noninvasively monitoring gene expression and plant transformation. Hortic. Res., 7.

Kshetry, A.O., Ghose, K., Alok, A., Devkar, V., Raman, V., Stupar, R.M., et al. (2025) A synthetic transcription cascade enables direct in planta shoot regeneration for transgenesis and gene editing in multiple plants. Mol. Plant, 18, 2066–2081.

Song, G.Q. and Sink, K.C. (2004) Agrobacterium tumefaciens-mediated transformation of blueberry (Vaccinium corymbosum L.). Plant Cell Rep., 23, 475–484.

Wang, G., Liu, M., Liu, G., Bao, Z., and Ma, F. (2022) Establishment and optimization of agrobacterium-mediated transformation in blueberry (Vaccinium species). Sci. Hortic., 304.

Yan, H., Ma, D., Yi, P., Sun, G., Chen, X., Yi, Y., and Huang, X. (2023) Highly efficient Agrobacterium rhizogenes-mediated transformation for functional analysis in woodland strawberry. Plant Methods, 19.

