## Supplementary Tables for "A robust and high-efficiency *Rhizobium rhizogenes* hairy root transformation platform for *Vaccinium*"

**Table S1.** Primers used in this study

| **Targeted gene** | **Primer Name** | **Sequence** | **Amplicon size (bp)** |
| --- | --- | --- | --- |
| *RUBY* | RUBY_F | CTATCCTTCGCAAGACCCTTC | 722 |
|  | RUBY_R | GGCTTGCCTATATCTTCCATG |  |
| *GFP* | GFP_1_F | ATACTCCAATTGGCGATGGCCCTG | 450 |
|  | M13PUC_R1 | CCCAGTCACGACGTTGTAAAACG |  |
|  | EGFP_qPCR_F | TTGCCGTCCTCCTTGAAGTC | 103 |
|  | EGFP_qPCR_R | TCTCGTTGGGGTCTTTGCTC |  |
| *WIND1* | WIND1_F2 | AGACTGGATCTGGTGGATCT | 534 |
|  | 35S_R2 | TGCGATAAAGGAAAGGCTATCA |  |
|  | WIND_qPCR_F | CGGTTCAGGTGTTCCTTCGA | 123 |
|  | WIND_qPCR_R | AAAGTCCCAAGCCAGAGACG |  |
| *ipt* | ipt_F1 | CCGTGGGCCTCATAATTGTA | 456 |
|  | ESR1_R1 | AAGCATTTATACTCTTCGCCAT |  |
|  | IPT_qPCR_F | TTGCACAGGAAAGACGTCGA | 146 |
|  | IPT_qPCR_R | GACGGGTCGTTCCTTTCAGT |  |
| *VcGAPDH* | VcGAPDH_F | CCGGAGCTGAGTTTGTTGTT | 309 |
|  | VcGAPDH_R | GACCACCTTCTTYGCACCAC |  |

**Table S2.** Antibiotic treatment combinations used for bacterial overgrowth control experiment

| **Antibiotic concentration (mg/L)** | **T1** | **T2** | **T3** | **T4** | **T5** | **T6** | **T7** | **T8** | **T9** | **T10** | **T11** |
| --- | --- | --- | --- | --- | --- | --- | --- | --- | --- | --- | --- |
| Cefotaxime | 300 | 200 |  |  | 200 | 200 |  | 100 | 300 | 300 | 0 |
| Timentin |  | 200 | 300 |  | 100 |  | 200 | 200 | 100 |  | 0 |
| Carbenicillin |  |  |  | 300 |  | 100 | 100 |  |  | 100 | 0 |

**Table S3.** *Vaccinium* genotypes used in this study

| **Genotype** | **Species** | **Section** |
| --- | --- | --- |
| Albus | *V. corymbosum* | *Cyanococcus* |
| Colossus | *V. corymbosum* | *Cyanococcus* |
| Optimus | *V. corymbosum* | *Cyanococcus* |
| Sharper | *V. corymbosum* | *Cyanococcus* |
| Elliottii | *V. elliottii* | *Cyanococcus* |
| AP3 | *V. stamineum* | *Polycodium* |

**Table S4.** Transformation efficiencies of six *R. rhizogenes* strains across *Vaccinium* genotypes

| **Strain**  **Genotype** | **AP3** | **Albus** | **Colossus** | **Elliottii** | **Optimus** | **Sharper** |
| --- | --- | --- | --- | --- | --- | --- |
| Ar.1193 | 3.5 ± 1.7 a | 10.8 ± 3.7 a | 6 ± 2.4 a | 1.3 ± 0 a | 3.4 ± 1.4 a | 6 ± 2.5 a |
| Ar. A4 | 5.8 ± 2.7 a | 71.7 ± 8 b | 85.2 ± 5.1 c | 1.9 ± 0 a | 43.7 ± 8.2 b | 29.6 ± 9.2 b |
| ATCC15834 | 4.4 ± 2.4 a | 53.7 ± 9 b | 44.6 ± 9 b | 1.3 ± 0 a | 10.5 ± 3.8 a | 12.9 ± 5 ab |
| C58C1 | 3.5 ± 1.7 a | 8.1 ± 2.8 a | 3.85 ± 1.6 a | 1.3 ± 0 a | 3 ± 1.2 a | 5.3 ± 2.2 a |
| K599 | 3.5 ± 1.7 a | 16.5 ± 5.4 a | 7.2 ± 2.9 a | 1.3 ± 0 a | 6 ± 2.4 a | 8 ± 3.3 ab |
| MSU440 | 3.5 ± 1.7 a | 12.5 ± 4.2 a | 4.5 ± 1.9 a | 1.3 ± 0 a | 4 ± 1.6 a | 6.9 ± 2.9 a |

Data is represented as Percentage transformation efficiency ± Standard error. Different letters indicate statistically significant differences among different treatments within the cultivars.
