## Supplementary Figures for "A robust and high-efficiency *Rhizobium rhizogenes* hairy root transformation platform for *Vaccinium*"

**Supplementary Figure 1.** **PCR-based confirmation of RUBY integration in induced hairy roots.** **a.** PCR-based molecular screening confirmed the presence of the *RUBY* gene (amplicon size: 722 bp) depicted with their corresponding RUBY-expressing roots below. Controls include + (plasmid DNA), WT (wild-type roots from tissue culture plants), and – (water control). Lanes 1–8 represent individual root samples. ‘Albus’ leaf and stem explants under

**a.** PCR-based molecular screening confirmed the presence of the *RUBY* gene (amplicon size: 722 bp) depicted with their corresponding RUBY-expressing roots below. Controls include + (plasmid DNA), WT (wild-type roots from tissue culture plants), and – (water control). Lanes 1–8 represent individual root samples. ‘Albus’ leaf and stem explants under **b**. Control leaf explant in half WPM media. **c**. Stem explant after infection with either Ar. A4 or K599.
