## Supplementary Files for "A robust and high-efficiency *Rhizobium rhizogenes* hairy root transformation platform for *Vaccinium*"

**Materials and Methods**

**Plant material and explants used in the study**

The southern highbush blueberry (SHB) cultivar ‘Albus’ was used for an initial assessment. Newly developed shoots were harvested from the field and surface disinfected using 10% commercial bleach solution for 10 min under agitation. The tissue was rinsed three times with sterile distilled water under the flow hood, and further excised to one node explants and planted in Half-strength Woody Plant Medium (HWPM) [1/2 WPM; (Lloyd and Mccown, 1980)+ ½ MS; (Murashige and Skoog, 1962)] with MS vitamins, 2 mg/L trans-Zeatin (tZ), 2% sucrose, and solidified with 6.5 g/L agar, pH 5.2. Clean plant material was further propagated through two node micro cutting. Two-month-old plants used for *R. rhizogenes*-mediated transformation were initially propagated and maintained in the Blueberry Breeding and Genomics Laboratory at University of Florida on MWPM. Plantlets were grown under a 16 h light/8 h dark cycle and maintained at 25°C until used.

**Vector construct information**

The *35S:RUBY* (named hereafter *RUBY*) plasmid used in this study was generated by He et al. (2020) and obtained from Addgene (plasmid #160908). This vector encodes a synthetic open reading frame that enables betalain biosynthesis, resulting in a vivid red pigmentation, and is driven by the Cauliflower Mosaic Virus (CaMV) 35S promoter. To generate a *ΔRUBY* negative control vector, the RUBY CDS was excised using the In-Fusion HD Cloning Kit (Takara Bio).

***R. rhizogenes-*mediated transformation**

The *RUBY* plasmid was introduced into *R. rhizogenes* strains K599 and Ar. A4 (Lifeasible) via electroporation. Transformed cells were cultured at 28°C in Tryptone Yeast extract (TY) medium supplemented with spectinomycin (50 µg/ml) and either streptomycin (50 µg/ml) or kanamycin (50 µg/ml), depending on the strain used. Bacterial cultures were grown overnight in TY liquid medium at 28°C, harvested by centrifugation at 4,000 rpm for 15 min, and resuspended in infection media (HWPM containing 100 µM acetosyringone [ACS]), adjusted to an OD₆₀₀ of 0.5–0.6. Leaf and stem explants were excised and subjected to two types of wounding: (1) a basal cut or (2) mechanically punctured using a sterile needle, then incubated in the infection medium for 10 minutes under continuous horizontal agitation (Fig. 1a).

Following infection, explants were transferred to co-cultivation medium (either half- or full-strength WPM, both containing 100 µM ACS) and incubated in the dark for four days. Plates were maintained in a growth chamber under the same conditions described previously. After co-cultivation, explants were rinsed with sterile water containing 300 mg/L cefotaxime to eliminate residual bacteria. As control, additional explants were co-cultivated using *R. rhizogenes* with the Δ*RUBY* vector*,* following the same protocol.

To evaluate optimal plant culture conditions, explants were transferred to either half- or full-strength WPM medium supplemented with 300 mg/L cefotaxime. Cultures were maintained in a growth chamber under the same conditions as mentioned earlier.

**Evaluation of six *Rhizobium* strains and optimization of antibiotic concentrations**

Based on the preliminary assessment, the cultivar ‘Albus’ was evaluated using four additional *Rhizobium* strains, Ar.1193, C58C1, ATCC15834, and MSU440 (Lifeasible), all transformed with the same reporter plasmid *RUBY.* The previous 2 strains were also included, resulting in a total of 6 strains in the study. The goal was to identify the best *Rhizobium* strain for transformation in ‘Albus’ and also determine the optimum antibiotic concentration and combination for each strain that could effectively control bacterial overgrowth. Three different antibiotics, cefotaxime, timentin, and carbenicillin, were tested either individually or in combination, resulting in a total of 11 different treatments including control (Table S2). The study utilized the best media composition and wounding method from the previous experiment for optimal results.

**Assessment of *Rhizobium* performance across *Vaccinium* germplasm**

To further assess the performance of *Rhizobium* strains across diverse germplasms in *Vaccinium*, all six strains were evaluated in species representing two taxonomic sections, maintained in the University of Florida Blueberry Breeding and Genomics Program (UF-BBGP). The Cyanococcus section included *V. corymbosum* cultivars ‘Colossus’, ‘Optimus’, ‘Sharper’, and ‘Falcon’, as well as *V. elliottii*. The Polycodium section, was represented by *V. stamineum* (‘AP3’) (Table S3). The plants were propagated and maintained as mentioned earlier and explants were collected from 2-month-old plants. This evaluation was conducted to compare strain effectiveness and plant response across genetically distinct *Vaccinium* backgrounds. The optimal antibiotic concentrations identified in the previous experiment were applied in this study.

***RUBY*-characterization in hairy roots**

Hairy roots expressing *RUBY* were visually scored on explants at 15, 25, 40, and 60 days post co-cultivation (DPCC). Images were captured using a Leica M165 FC stereo fluorescence microscope and processed with Leica Application Suite X software. Genomic DNA was extracted from hairy roots collected between 50 and 60 DPCC using the CTAB method (Doyle, 1991). PCR amplification was performed with *RUBY*-specific primers (Table S1) using the GoTaq® Green Master Mix (Promega). The PCR protocol included an initial denaturation at 95°C for 5 minutes; 35 cycles of 95°C for 30 seconds, 60°C for 45 seconds, and 72°C for 45 seconds; followed by a final extension at 72°C for 7 minutes. Amplicons were observed on a 1% agarose gel stained with SYBR Safe DNA Gel Stain (Thermo Fisher Scientific).

**Plant regeneration following hairy root transformation**

Plant regeneration was attempted through two distinct approaches. In the first approach, the

*RUBY*-expressing roots were transferred to callus induction media (HWPM supplemented with 2 mg/L of trans-Zeatin and 2 mg/L of 2,4-Dichlorophenoxyacetic acid). This step was performed to evaluate the *RUBY* expression in callus tissue and induce somatic embryos as a preliminary approach for regeneration, ultimately facilitating shoot development. In the second approach, the most effective *Rhizobium* strain was selected and transformed with the plasmid *pESR1::ipt::35S::WIND1::YLCV::GFP*, hereafter referred to as *DRGFP*, a kind gift from Dr. Patil (Texas Tech University). The plasmid contains plant morphogenic regulators, *ipt* (*Isopentenyl transferase*) and *WIND1* (*WOUND INDUCED DEDIFFERENTIATION1*), which are involved in plant wound repair and regeneration, *GFP* (*Green Fluorescent Protein*) as a marker gene for easy phenotyping of transformed cells and *bar* (*Bialaphos resistance*) gene as plant selection marker (Kshetry *et al.*, 2025). The cultivar ‘Albus’ was chosen for transformation, and the plants were subcultured every three weeks under optimal media and antibiotic conditions determined in the above experiments.

**Screening of regenerated shoots**

The shoots that regenerated following hairy root transformation using the above approaches were visually screened for the presence of *GFP* using the fluorescent stereomicroscope, LEICA M165 FC. Further confirmation of transgene integration was performed by PCR using primers to *GFP*, *WIND1*, *ipt* and VcGAPDH (Table S1). PCR conditions were identical to those described for *RUBY*, and amplification products were resolved on a 1% agarose gel as described previously.

**Data analysis**

Data was collected from five replicates, each consisting of 30 explants. Hairy root induction efficiency was calculated as the proportion of explants that developed hairy roots for each wound type, relative to the total number of explants. At each time point, mean rooting efficiencies between cut and puncture treatments were compared using Student’s t-test.

Transformation efficiency was estimated by dividing the number of explants producing *RUBY*-expressing hairy roots by the total number of explants per treatment. Given the high frequency of zero counts, a zero-inflated negative binomial (ZINB) mixed-effects model was implemented using the ‘*glmmTMB’* package in R (R Core Team, 2024). Bacterial strain, explant type, culture medium, and DPCC were treated as fixed effects, while replicates were included as random effects.

**References**

Doyle, J. (1991) DNA Protocols for Plants. In: Molecular Techniques in Taxonomy (Hewitt,G.M., Johnston,A.W.B., and Young,J.P.W., eds) , pp. 283–293. Berlin, Heidelberg: Springer Berlin Heidelberg.

He, Y., Zhang, T., Sun, H., Zhan, H., and Zhao, Y. (2020) A reporter for noninvasively monitoring gene expression and plant transformation. Hortic. Res., 7.

Kshetry, A.O., Ghose, K., Alok, A., Devkar, V., Raman, V., Stupar, R.M., et al. (2025) A synthetic transcription cascade enables direct in planta shoot regeneration for transgenesis and gene editing in multiple plants. Mol. Plant, 18, 2066–2081.

Lloyd, G. and Mccown, B.H. (1980) Commercially-feasible micropropagation of mountain laurel, Kalmia latifolia, by use of shoot-tip culture.

Murashige, T. and Skoog, F. (1962) A Revised Medium for Rapid Growth and Bio Assays with Tobacco Tissue Cultures. Physiol. Plant., 15, 473–497.

R Core Team (2024) R: A Language and Environment for Statistical  Computing.
